# Amyloid-*β* peptide dimers undergo a random coil to *β*-sheet transition in the aqueous phase but not at the neuronal membrane

**DOI:** 10.1101/2020.12.31.424964

**Authors:** Hebah Fatafta, Mohammed Khaled, Abdallah Sayyed-Ahmad, Birgit Strodel

**Affiliations:** Institute of Biological Information Processing (IBI-7: Structural Biochemistry), Forschungszentrum Jülich, 52425 Jülich, Germany; Department of Physics, Birzeit University, PO Box 14, Birzeit, Palestine; Institute of Theoretical and Computational Chemistry, Heinrich Heine University Düsseldorf, 40225 Düsseldorf, Germany

**Keywords:** amyloid-*β*, oligomers, Alzheimer’s disease, MD simulations, transition networks

## Abstract

The aggregation of amyloid *β*-peptides into neurotoxic oligomers is a key feature in the development of Alzheimer’s disease. Mounting evidence suggests that the neuronal cell membrane is the main site of oligomer-mediated neuronal toxicity. To gain a detailed understanding of the mutual effects of amyloid-*β* oligomers and the neuronal membrane, we carried out a total of 12 *µ*s all-atom molecular dynamics (MD) simulations of the dimerization of the full-length A*β*42 peptide in the presence of a lipid bilayer mimicking the *in vivo* composition of neuronal membranes. The conformational changes of A*β*42 resulting from its dimerization and interactions with the neuronal membrane are compared to those occurring upon its dimerization in the aqueous phase, which is also tested by 12 *µ*s of MD simulations. We find that the interactions with the neuronal membrane decrease the order of the A*β*42 dimer by attenuating its propensity to form a *β*-sheet structure. The main lipid interaction partners of A*β*42 are the surface-exposed sugar groups of the gangliosides GM1. A*β*42 dimerization in solution, on the other hand, is characterized by a random coil to *β*-sheet transition that seems to be on-pathway to amyloid aggregation. As the neurotoxic activity of amyloid oligomers increases with oligomer order, the results suggest that GM1 is neuroprotective against A*β*-mediated toxicity by inhibiting the formation of ordered amyloid oligomers.

## 1 Introduction

The aggregation of proteins into amyloid deposits is a notable feature of multiple diseases. In Alzheimer’s disease (AD), it is the amyloid *β*-peptide (A*β*) that aggregates and subsequently accumulates within the neural tissue [1, 2, 3]. A*β* has two C-terminal variants (A*β*1-40 and A*β*1-42) that are naturally produced in the brain by the cleavage of the amyloid precursor protein [4]. The A*β*42 variant is known to be more toxic and shows a higher aggregation propensity, ranging from the formation of dimers to dodecamers [5] and ultimately amyloid fibrils [6]. An increasing number of studies suggest that the soluble oligomers formed in the earlier stages of the aggregation process, and not the large insoluble fibrils, are the main cytotoxic species affecting the severity and progression of AD [7, 8, 9]. Therefore, in the past one to two decades a shift has been made toward exploring the assembly, structure, and toxicity of these oligomers rather than fibrils [10, 11]. A*β* dimers have been reported to be the smallest toxic oligomer affecting synaptic plasticity and cause memory impairment [12, 13]. In addition, several experimental studies postulate that A*β* dimers might act as a nucleation seed for further aggregation [14, 15, 16]. Therefore, a detailed characterization of A*β* dimerization is an essential step toward a better understanding of the aggregation process. However, it is challenging to explore the dimer structure experimentally due to the transient nature resulting from the A*β* aggregation tendency, its plasticity, and the existence of the dimer in equilibrium with the monomer and higher order oligomers [17, 18]. Molecular dynamics (MD) simulations are able to close this gap in knowledge by providing information about the dimer structure at the atomic level and its temporal evolution [19, 20, 21, 22, 23, 24, 25, 26, 27]. However, the previous simulations of A*β* dimers modeled them in the aqueous phase only, thus lacking the detailed cellular context. Consideration of the latter is necessary if one wishes to reveal the mechanism of toxicity which is believed to occur via damaging the lipid membrane of neurons by A*β* oligomers [28].

The interactions of A*β* with lipid membranes affect its aggregation pathways from soluble monomers into toxic oligomers or fibrils [29]. Importantly, different lipids of the neuronal membrane have been observed to have different effects on the aggregation of the peptide. For example, the liquid-ordered phases of lipid membranes rich in sphingomyelin (SM) and cholesterol (CHOL) were found to enhance A*β* aggregation [30], while monosialotetrahexosylganglioside (GM1) was found to both accelerate A*β* aggregation as well as inhibit it depending on its density and relative abundance [31]. In a previous MD study, we explored the effect of the membrane composition on monomeric A*β*42 and showed that physiologically relevant levels of SM, but not GM1, induce a *β*-sheet-rich structure in A*β*42 monomers [32]. Moreover, in another simulation study we observed a reduction in *β*-sheet formation upon interaction with GM1 [33]. The generalized picture from these and other simulation studies [34, 35] points to the role of electrostatic interactions between A*β* and the lipid hydrophilic groups in driving adsorption of the peptide to the membrane, whereas the interactions of the mostly hydrophobic C-terminal residues with the hydrophobic core drives the membrane insertion of A*β*. However, the latter has not been observed yet by simulations – unless A*β* preinserted into a membrane was simulated [36, 37, 38, 38, 39, 40] – but was recorded by experimental means for both monomeric [41] and aggregated [42] A*β*. However, it is not known yet what the structure of membrane-inserted A*β* aggregates is. Membrane-inserted monomeric A*β* adopts a helical structure [41, 43] and insertion inhibits the fibrillation [43] but leads to A*β* aggregates with ion channel-like structures [42].

In this study, we use an aggregate of 24 *µ*s MD simulations to investigate the dimerization of the full-length A*β*42 peptide both in solution and in the presence of a model lipid bilayer including six lipid types to mimick that of a neuronal cell membrane: 38% 1-palmitoyl-2-oleoyl-sn-glycero-3-phosphocholine (POPC), 24% 1-palmitoyl-2-oleoylsn-glycero-3-phosphoethanolamine (POPE), 5% 1-palmitoyl-2-oleoyl-sn-glycero-3-phospho-L-serine (POPS), 20% CHOL, 9% SM, and 4% GM1 (Figure 1). To the best of our knowledge, this simulation study breaks new ground at two fronts: i) it exceeds the simulation time of previous studies modeling A*β*–membrane interactions by an order of magnitude; ii) the aggregation of A*β* modified by lipid bilayers containing more than three different lipid types has not been studied yet using MD simulations. Most of the previous MD studies examining the effects of a membrane on A*β* dimerization considered simpler lipid bilayers [44, 45, 46]. Lipid bilayers of comparable complexity as the one modeled here were thus far only simulated at the coarse-grained level [47, 48]. Another highlight of our study is the analysis of the aggregation pathways by transition networks [49, 50, 51], which excels at elucidating the similarities and differences between A*β* dimerization in solution and at the neuronal membrane. A main finding is that the presence of the neuronal membrane, on the one hand, reduces the dynamics of membrane-bound A*β*42 while, at the same time, it inhibits *β*-sheet formation. Here, a key player are the sugar groups of GM1, which form hydrogen bonds with the peptide, thereby reducing the possibilities for intra- and inter-peptide hydrogen bonds. In contrast, the dimerization in the aqueous phase is characterized by a random coil to *β*-sheet transition, leading to a *β*-sheet structure similar to the one found in some of the A*β* fibrils.

**Figure 1:**
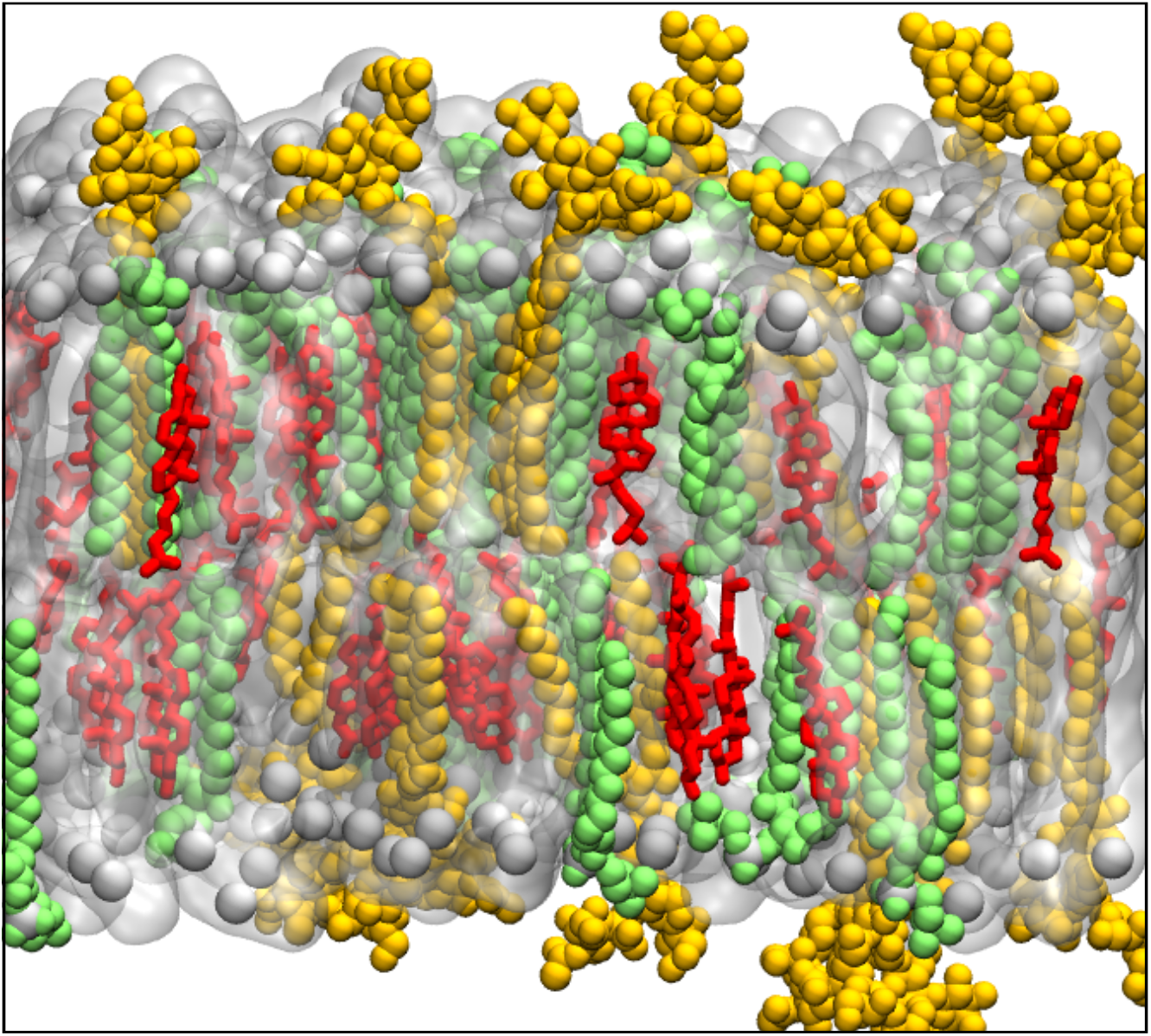
A snapshot of the neuronal membrane containing 38% POPC, 24% POPE, 5% POPS (collectively shown as silver surface with their phosphorous atoms indicated by silver spheres), 20% CHOL (red sticks), 9% SM (green spheres), and 4% GM1 (yellow spheres).

## 2 Results and Discussion

### 2.1 Characteristics of the neuronal membrane

Before we studied the interaction of A*β* with the neuronal membrane, we equilibrated the membrane without peptides being present for 1 *µ*s. Afterwards, 6 × 2 *µ*s MD simulations with two A*β*42 peptides added to the system were performed. These simulations are used to characterize the neuronal membrane and its interaction with A*β*42.

#### Mass density along the bilayer normal

The mass density profile of the different lipids and water along the membrane *z*-axis (Figure S1) shows the characteristic features of a lipid bilayer and provides information about the bilayer thickness. The positions of the headgroups peak at similar locations for POPC, POPE, POPS, and SM. CHOL, on the other hand, prefers locations shifted toward the hydrophobic core of the bilayer, while GM1 is farther away from the bilayer center resulting from the the sugar groups sticking out of the membrane (Figure 1). The headgroup–headgroup distance of POPC, POPE, and POPS gives an indication for the bilayer thickness, which was determined as 4.65 ± 0.03 nm.

#### Lipid clustering

The radial distribution function (RDF) of for all possible lipid pairings was calculated to monitor lipid clustering resulting from lipid–lipid interactions (Figures 2 and S2 in the Supporting Information). For the calculation of these pair correlation functions, the phosphorous atoms POPC, POPE, POPS, and SM as well as the oxygen atoms of CHOL and GM1 were considered and the corresponding RDFs were averaged over the upper and lower leaflet.

**Figure 2:**
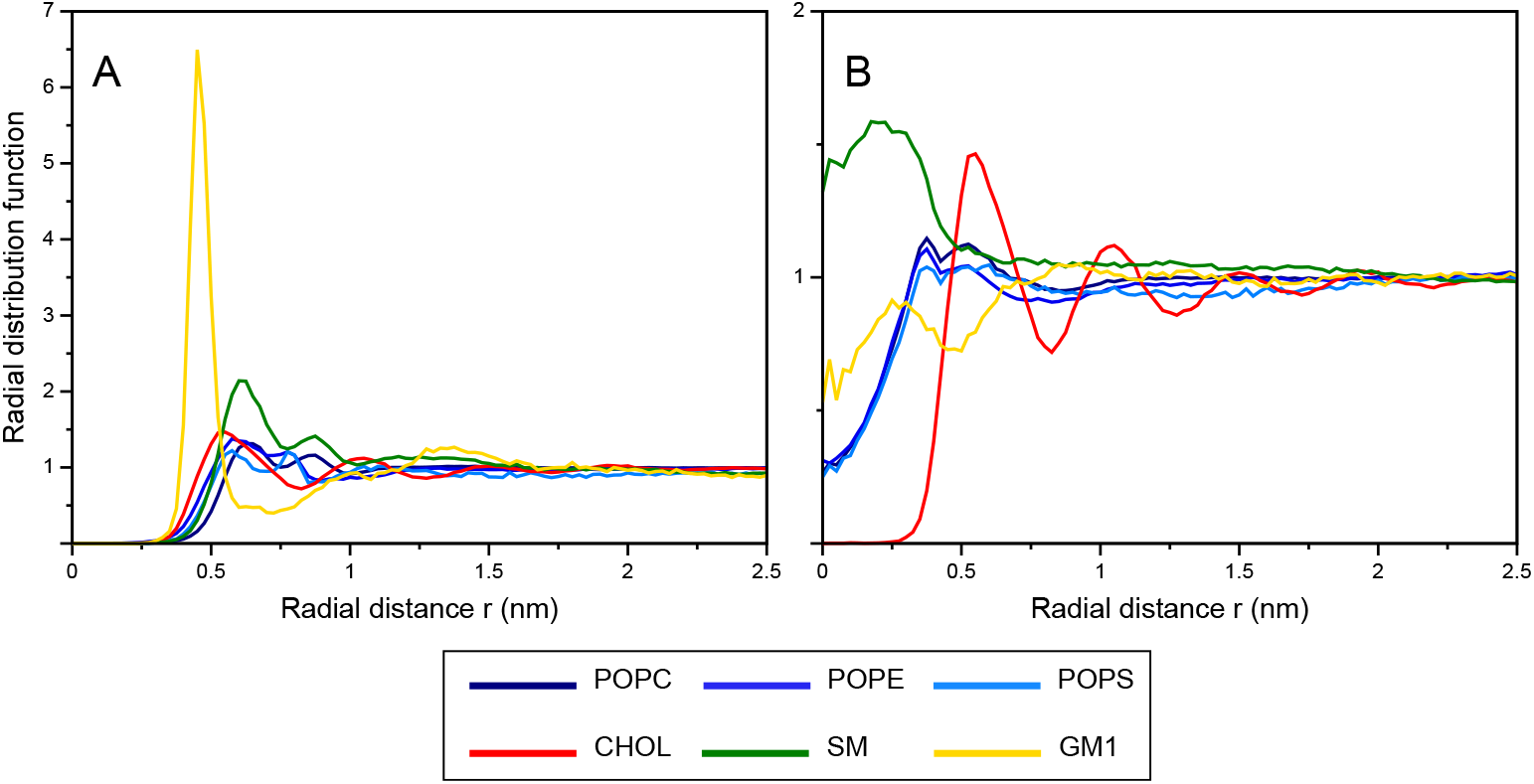
Radial distribution functions for (A) lipid pairings of identical type, (B) lipid–CHOL pairings. The P atoms of POPC, POPE, POPS, and SM and the O atoms of CHOL and GM1 were used for the RDF calculations. The *x*-axes show the distances between the respective atom pairs. The colors of the graphs refer to the lipids as indicated in the color key below the plots. Pairs with RDF > 1 are considered to cluster. The plots for the RDFs of the remaining mixed lipid pairs are shown in Figure S2.

The first sharp RDF peak indicating self-clustering of lipids is seen at 0.6 nm for POPC, POPE, POPS, CHOL, and SM and at 0.45 nm for GM1. The GM1–GM1 clustering is considerably stronger than those for the other lipids. Thus, taking the low proportion of GM1 in the membrane (4%) into account, one can conclude that GM1 has a strong tendency to self-assemble leading to lipid sorting. Clustering of mixed lipid pairs is observed for combinations of POPC, POPE, and POPS with each other at 0.6 nm. Interestingly, the RDF of POPE–POPS has a higher peak compared to self-clustering of POPE and POPS, respectively. The avoidance of self-association of POPS is understandable given that it is a negatively charged lipid. SM and GM1 tend to cluster at 0.5 nm with POPC and POPE. CHOL prefers to cluster near SM as the corresponding RDF has a higher peak compared to the other RDFs involving CHOL. A notable feature of all CHOL–lipid RDFs is the fact that the RDFs do not approach zero at the origin. Cholesterol resides deeper inside the bilayer and is shorter compared to the flexible tails of the lipid. Hence, it is possible that the oxygen atom of CHOL and the phosphorous atoms of the other lipids or the the O atom of GM1 used for calculating the RDFs are above each other. In most cases, the RDFs for lipid pairs showing clustering are characterized by a single sharp peak at 0.6 nm, indicating lateral short-range ordering or even lipid sorting. More peaks appear at larger distance, manifesting an increase in the ordering up to 1.0 nm. At distances beyond 1.0 nm, all RDFs approach unity, highlighting the equilibration of different lipids. Similar lipid clusterings have been reported by studies on bilayer models resembling the mammalian plasma membrane [39, 52] and in multiple MD studies of simpler bilayer models that investigated CHOL mixing with sphingomyelin and glycerophospholipids lipids [53] or in different phospholipid membranes [54, 55].

#### Hydrogen bonding between lipids

To elucidate the lipid–lipid interactions underlying the RDFs, the average numbers of hydrogen bonds (H-bonds) formed between the different lipid pairs over simulation time were evaluated (Figure S3). Hydrogen bonds are crucial for structure formation in and between biomolecules, such as packing of lipids in membrane [56]. Figure S3 shows that H-bonds formed between the hydroxyl of CHOL and the amide group of SM. This not only explains the clustering tendency between SM and CHOL, but also the buried location of CHOL within the lipid bilayer (Figure S1). The propensity of SM to form H-bonds with itself also gives rise to its self-clustering. Similarly, the sorting GM1 results from its ability to build a network of H-bonds via its sugar headgroups, despite it being negatively charged. However, the negative charge of both GM1 and POPS explains why these two lipids do not co-cluster. The self-clustering of POPC and POPE is expected due to their dominance in the lipid bilayer (38 and 24%, respectively), which makes it more likely for these molecules to be found in each other’s neighborhood. However, POPE possesses an amine group providing a site for additional intermolecular hydrogen bonding. This explains the sharp RDF peak observed for POPE self-clustering as compared to POPC–POPC clustering and the co-location of POPE and POPS. Our findings agree with those of previous MD studies that highlighted the importance of H-bonds in stabilizing lipid clusters [53, 57]. However, the role of such H-bonds has also been debated [58]. In particular, some MD and experimental studies point to the effect of the lipid head group properties allowing their interaction with the neighboring lipids. For example, they relate the clustering of SM with CHOL to the SM headgroup, which has the ability to orient toward the lipid bilayer providing CHOL with better shielding from the solvent, thus making the SM–CHOL interactions more favorable [59, 60, 61, 62, 63]. Despite these differences, it is generally accepted that H-bonds contribute to stabilizing lipid clustering.

#### Acyl chain ordering

We calculated the deuterium order parameter, *S*_CH_, of the C–H bonds in the lipid tails for all lipids (Figure 3) to gain a more detailed picture about the structure of the acyl chains. They all have the typical trend seen in previous experimental and MD studies [64, 65]. However, a slight increase in the order parameter values are seen in our MD simulations. This might be due to the effect of having different lipid types in the membrane, including cholesterol and sphingomyelin that are known for their role in increasing lipid ordering [58, 66]. Notably, we find the chains of GM1 and SM to be the most ordered. A closer look reflects how the palmitoyl tails of the different lipids reach a plateau at 0.35–0.4 for the order parameters of carbon atoms 4–10, which then decrease to zero at the ends of the tails, while the olyoel tails exhibit a strong drop in order at the double bond positioned at carbon atom 10. It has been reported that the orientation of the double bond (C=C) with respect to the bilayer normal might cause such a drop in the order parameter [67, 68]. However, no reduction in *S*_CH_ at the double bond position of SM (carbon atoms 4–5) was detected, which agrees to other MD studies [69] but not to the one by Niemalä et al. [70] who reported a reduced *S*_CH_ for the double bond of SM. Since there is a lack of experimental data in this regard, these conflicting findings cannot be explained.

**Figure 3:**
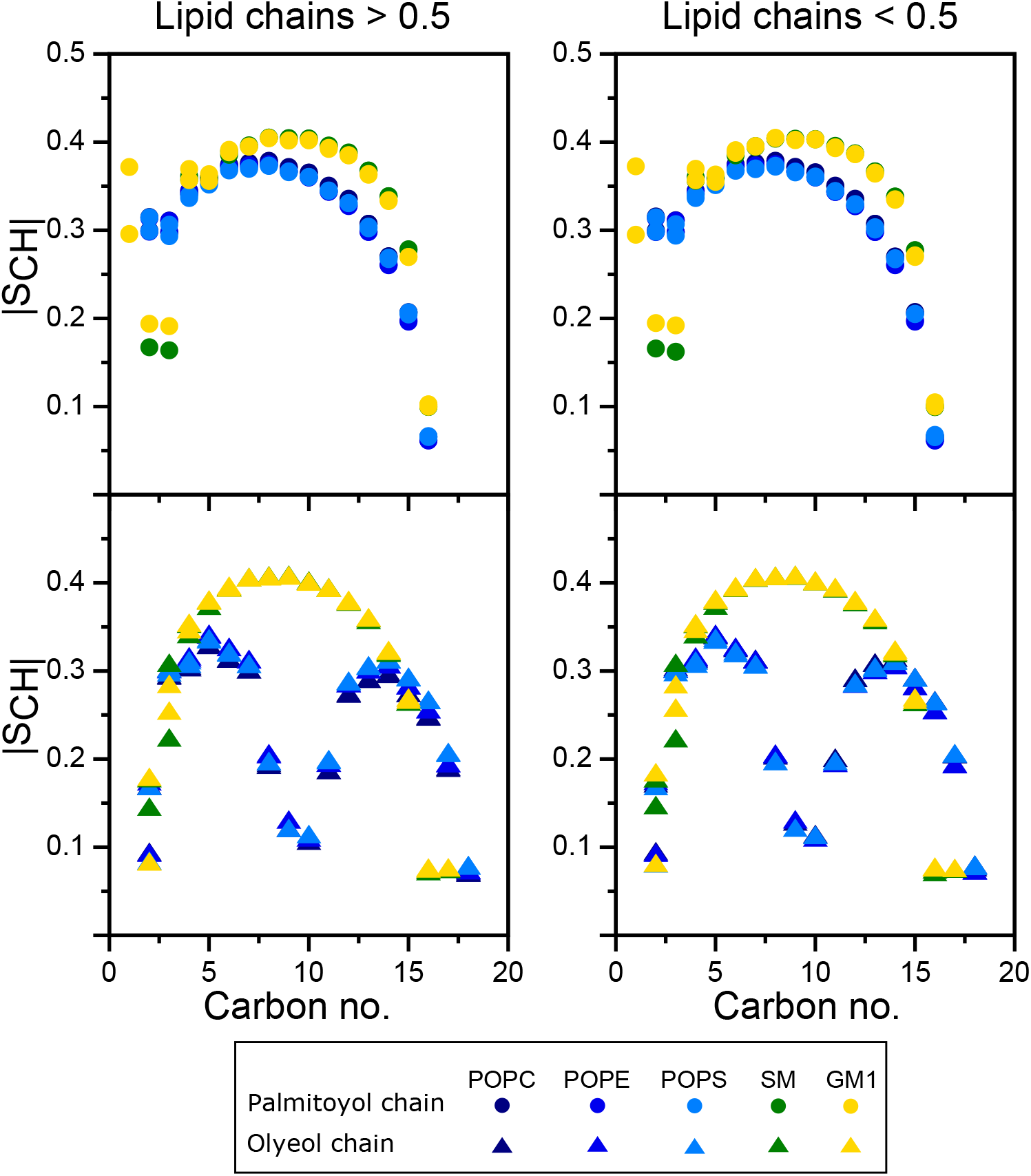
Average order parameters of the acyl chains (top: palmitoyol chains; bottom: olyeol chains) of each lipid component of the neuronal membrane, distinguishing between lipids that are more than 0.5 nm away from A*β*42 (left) and those within 0.5 nm of the peptide (right).

### 2.2 Interactions of A*β*42 with the neuronal membrane

To understand the effects of the neuronal membrane on the aggregation of A*β*42, we analyzed the 12 *µ*s of MD data in the presence of the lipid bilayer and compared the aggregation to that in the aqueous phase, for which another 6 × 2 *µ*s of MD simulations were performed. Before the aggregation process is discussed in detail, we assess whether and how the two peptides, which were initially placed above the membrane, interact with the latter.

#### Membrane adsorption

To follow the association between A*β*42 and the neuronal membrane, we calculated the minimum distance of both peptides from the lipid bilayer surface for each of the six simulations (Figure S4). It can be seen that peptide 1 usually interacts more closely with the membrane than peptide 2. Moreover, if one of the distances is » 0.5 nm, implying that the corresponding peptide is in solution, then very often this also the case for the other peptide, indicating that both peptides tend to be associated as a dimer with the membrane. In MD run 1 dimerization occurred at around 500 ns on the membrane surface. Figure S5 shows representative snapshots for the membrane association of A*β*42, including one for loose attachment in panel A. Figure S5B represents the situation where peptide 1 is in close contact with the membrane, while peptide 2 is a bit further away. The less common opposite situation with peptide 2 being closer is depicted in Figure S5C, while Figure S5D represents the case where both peptides are in intimate contact with the membrane. Figure S5 further suggests that A*β*42 has a preference to interact with GM1 as compared to the other lipids, and that *β*-sheets are the dominating secondary structure in peptide 2 but not in the more membrane-bound peptide 1. The analysis of the contacts between A*β*42 and the various lipids confirms that the peptide has a high tendency to associate with GM1, followed by POPC, POPE, and POPS (Figure S6). Here, it should be emphasized that these contacts are not normalized but absolute values. Considering that only 4% of the lipids are GM1 while the phospholipids make up for more than two third of the membrane, one can thus conclude that A*β*42 is highly attracted by GM1. Interestingly, almost no contacts are formed with CHOL or SM.

#### Interaction energies and H-bonds

To rationalize the driving force that controls A*β*42 interaction with the lipid bilayer surface, the interaction energy of each A*β*42 residue with each of the components of the neuronal membrane was calculated and partitioned into its electrostatic (*E*_Coul_) and Lennard-Jones (*E*_LJ_) contributions (Figure 4). These two interaction types contribute differently to the different peptide–lipid interactions, showing a strong dependence on the lipid type. Notably, the lipid interactions of peptide 1 are energetically more favorable than those of peptide 2, agreeing to the observation that peptide 1 interacted more intimately with the membrane (Figure S6). Our results suggest that the major driving force for the membrane association of the peptides is the electrostatic attraction to POPC, POPE and POPS, especially via the considerably charged N-terminal region (D1 to Y10) and residues F20 to A30. Some charged residues from the N-terminal region are found to have the strongest interactions, like D1, E3, and D7 with POPE, D1 with POPC, and R5 and POPS. The latter interaction involves H-bond formation (Figure S7), which is enabled via the carboxylate group of POPS, while the primary ammonium group of POPE forms H-bonds with D1, E3, and D7. The tertiary ammonium group of POPC, on the other hand, does not allow for H-bond formation, leading to an only small H-bond propensity between POPC (via its phosphate group) and A*β*42. The interactions between GM1 and A*β*42 involve both Coulomb and Lennard-Jones energies (Figure 4) and are facilitated by the sugar headgroups of GM1, which stick out of the membrane and are therefore particularly accessible for A*β*42. Also the interactions with GM1 derive from a considerable number of H-bonds, which involve almost all residues of the two peptides, but especially of peptide 1.

**Figure 4:**
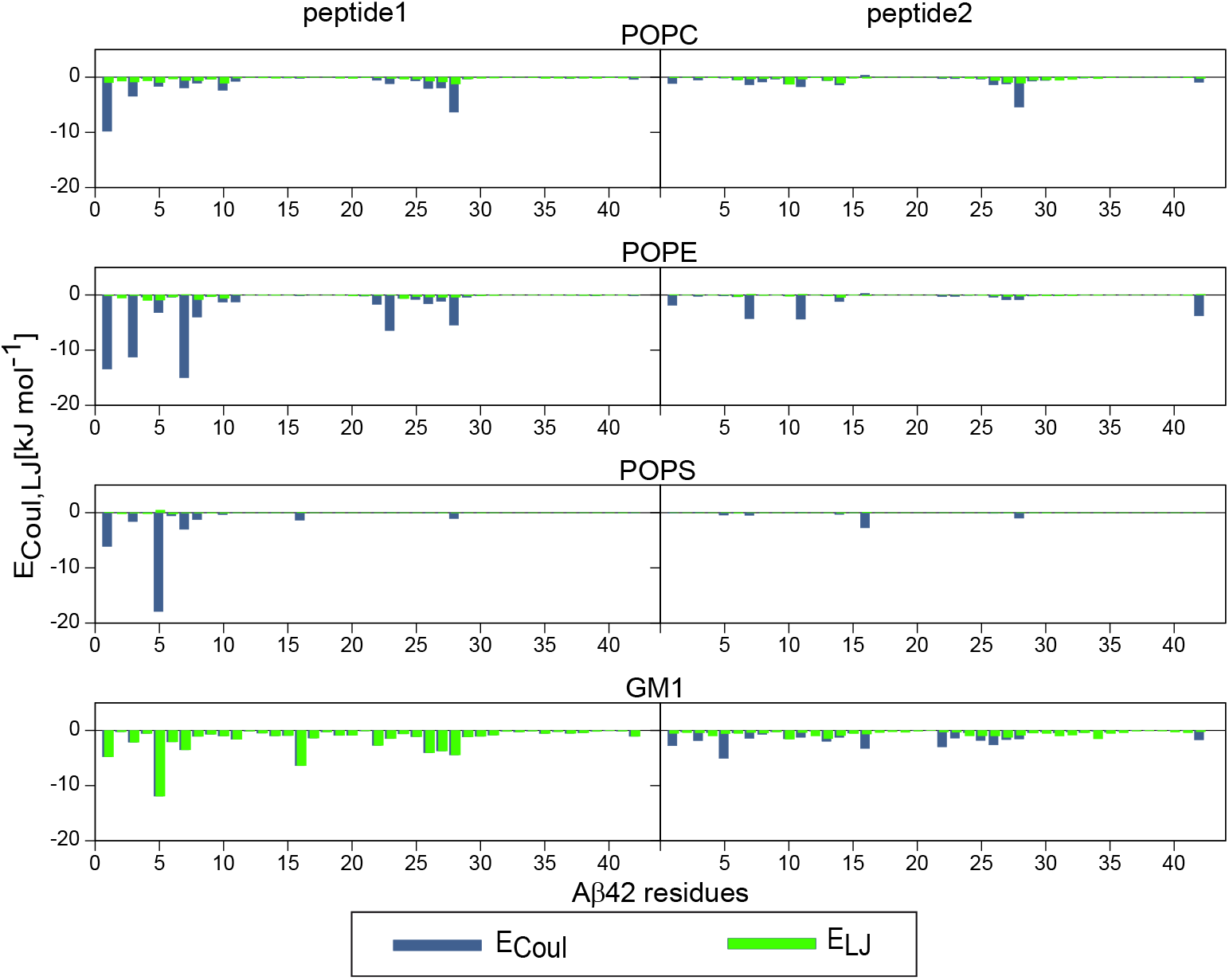
The average interaction energies of peptide 1 (left) and peptide 2 (right) with each of the components of the neuronal membrane. Electrostatic and Lennard-Jones energies are shown in blue and green, respectively.

Our observations are similar to those of previous studies of A*β* monomers interacting with lipid bilayers containing GM1 [32, 33, 45]. From a simulation study of an A*β*42 dimer interacting with a ternary membrane containing POPC, CHOL and GM1, Manna et al. found that charged residues preferentially interact with POPC, while polar and hydrophobic residues favor H-bond formation with GM1 sugar groups [45]. Moreover, a fluorescence-imaging study showed that A*β* prefers to colocalize with GM1-rich domains on cell membranes [71]. No direct interaction between CHOL contained in the neuronal membrane simulated here and A*β*42 was observed (Figure S6), which can be explained by the deeper, not surface-exposed position of CHOL inside the membrane. Since in our simulations no noteworthy membrane insertion of A*β*42 took place, contacts between the peptide(s) and CHOL were thus not established. This finding is in agreement with previous results by Fantini et al. [72], who further showed that CHOL induces a glycolipid conformation that is optimal for A*β* recognition via polar interaction with GM1 oligosaccharides as well as NH and OH of sphingosine. Interestingly, even though SM has the same headgroup as POPC, which is also loacated at a similar position along the bilayer normal (Figure S1), A*β*42 hardly interacted with SM in our simulations (Figure S6). This can be understood by considering interactions of SM with itself and the other lipids as reflected by the RDFs (Figures 2 and S2). It seems that SM prefers to interact with other lipids including itself, especially with saturated lipid chains and CHOL. These interactions are further enhanced by the formation of a hydrogen bond network at the membrane interface, thereby preventing it to interact with A*β*42.

#### Peptide effects on the neuronal membrane

Since both peptides interact with the neuronal membrane, a question is whether they have an effect on the lipid bilayer properties. To answer this question, we calculated the lipid order parameter for the lipids that are within 0.5 nm of the peptide when adsorbed to the membrane (Figure 3). The results suggest no notable change in the lipid order parameter due to the presence of A*β*42. The analysis of the bilayer thickness (Figure S8) indicates slight disturbances of the bilayer thickness originating from the membrane adsorption of A*β*42, leading to deviations of about ± 0.1 nm from its average value of 4.65 ± 0.03 nm. This observation correlates with the conclusion that the peptides interact only with the lipid head groups but do not insert into the membrane, thereby preventing larger changes in the membrane order and thickness.

### 2.3 Dimerization of A*β*42 in solution and at the neuronal membrane

#### Aggregation pathways

In order to unravel the aggregation pathways, we computed transition networks (TNs) for both the dimerization in the aqueous phase and in the presence of the neuronal membrane. This approach implies the characterization of the MD snapshots using beneficial descriptors. Here, we considered the aggregate state (monomer or dimer), the number of hydrophobic contacts between the peptides in a dimer, and the number of residues in *β*-strand conformation as descriptors. To further simplify the TNs, we grouped both the number of hydrophobic contacts and the number of residues in *β*-strand conformation in blocks of five such that we end up with ranges h1 to h12 and b1 to b6. For example, h1 and b1 stand for hydrophobic contacts and number of residues in *β*-strand conformation, respectively, ranging from 1 to 5. The maximum range h12 involves between 56 and 60 hydrophobic contacts and b6 means that between 26 and 30 residues per peptide adopted a *β*-strand conformation.

A marked difference between the resulting TNs (Figure 5) is the larger number of nodes and more transitions occurring between nodes for A*β*42 dimerization in solution. This implies that the neuronal membrane restricts the conformational diversity of the peptides. Irrespective of this difference, both TNs are characterized by two regions: the monomeric region (on the left side of the TNs) that evolves to the dimeric region (in the middle and right side of the TNs). These regions are connected by several bridging nodes, which, on average, are characterized by a higher amount of *β*-sheet (i.e., larger *n* in the descriptor b*n*) in the case of the solution system. For each TN, a representative bridging node is indicated by a green circle in Figure 5, (2, h2, b6) for the solution system and (2, h2, b2) for the membrane system, which are further augmented by a characteristic structure. In solution, there are more transitions between monomers and dimers visible, indicating a higher amount of association and dissociation events. A closer inspection of both TNs reveals how the two peptides evolve from the monomeric random coil state, which is presented by node (1, 0, 0) with no interpeptide hydrophobic contacts and no residues in *β*-strand conformation, to dimers with few hydrophobic contacts contained in the nodes (2, h1, b*n*), where b*n* ranges from b1 to b6 indicating a *β*-strand increase, by passing through nodes (1, 0, b*n*) and (2, 0, b*n*). The dimers with no hydrophobic contacts are so-called encounter complexes, where the minimal distance between the two monomers fell below 4.5 nm and which evolve into stable dimers by increasing their contact area leading to interpeptide contacts. This process of dimer stabilization is accompanied by an increase in *β*-strand amount. In solution, the dimers reach a higher amount of interpeptide hydrophobic contacts, reaching states (2, h12, b6) and (2, h13, b5) where between 50 and 70% of all A*β*42 residues are part of a *β*-sheet. In the presence of the neuronal membrane, both the hydrophobic contact area and *β*-sheet amount are overall smaller, with the maximal values being (2, h10, b4) and (2, h9, b5). In this case, some of the interpeptide contacts are replaced by peptide–lipid contacts, which in turn also affects the formation of *β*-sheets.

**Figure 5:**
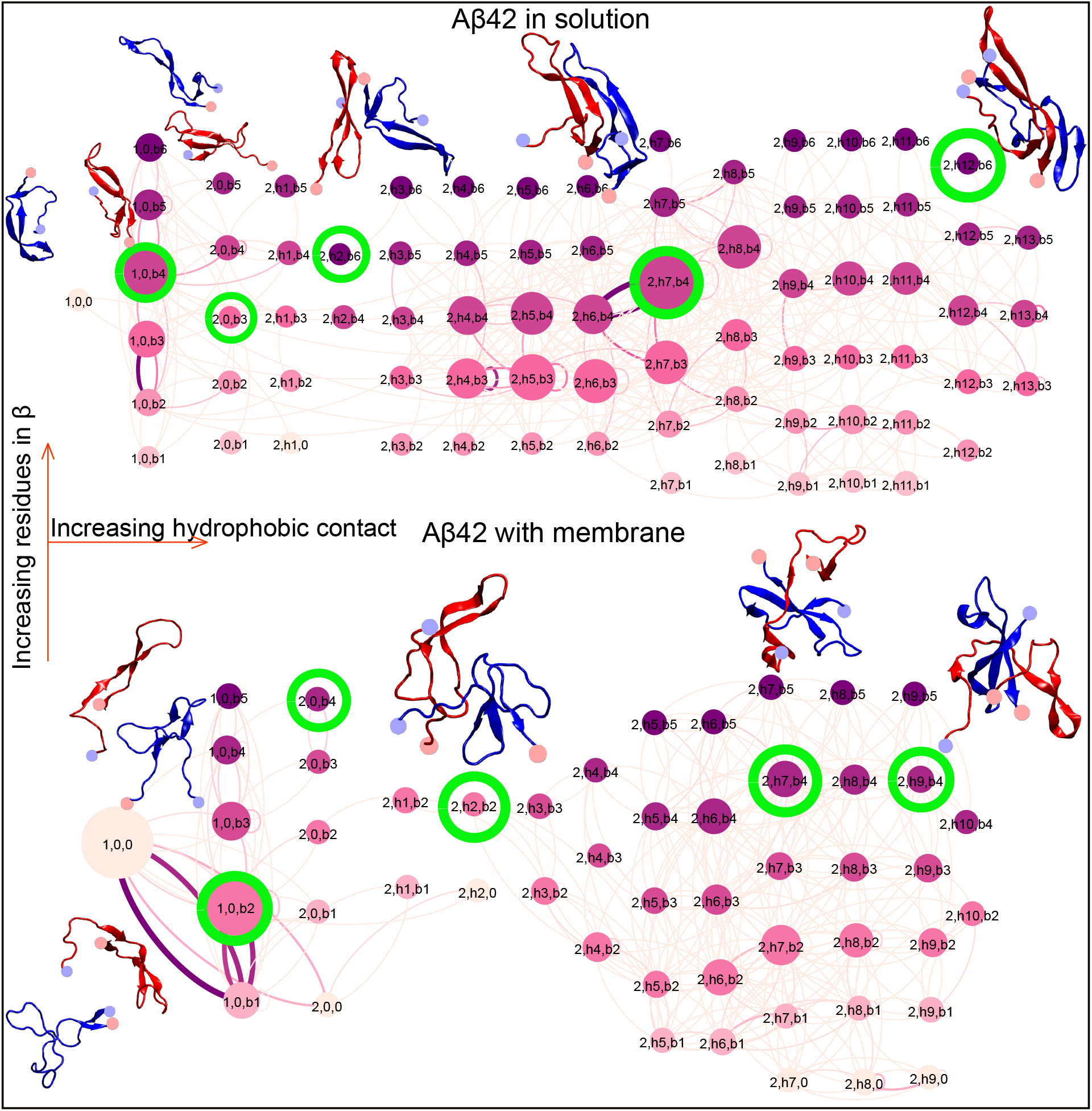
The TN for A*β*42 dimerization in the aqueous phase (top) and in the presence of the neuronal membrane (bottom). Each node is defined by three descriptors: oligomer size, number of interpeptide hydrophobic contacts, and number of residues in *β*-strand conformation. The last two descriptors are grouped in blocks of 5 and are named h1 to h12 for hydrophobic contacts and b1 to b6 for residues in *β*-strand conformation. For example, the node (2, h1, b1) represents a dimer that has hydrophobic contacts in the range 1–5 and has 1–5 residues in *β*-strand conformation. The nodes are connected by edges that represent transitions between the connected peptide states. The size of the nodes and the thickness of the edges are proportional to the respective probability. They are colored based on the descriptor reflecting the number of residues in *β*-strand conformation (from light pink for no *β*-sheets to dark purple for the maximum amount of *β*-sheets corresponding to the range b6). For the nodes circled in green representative peptide conformations are shown as cartoon, using red for peptide 1, blue for peptide 2, and spheres to indicate the N- and C-termini (light blue and light red, respectively).

This conclusion is confirmed by the representative structures shown along with the TNs (Figure 5) and the structures illustrating the membrane adsorption of the dimer (Figure S5). The membrane-adsorbed dimer structures are more compact than the dimer structures prevailing in solution. The latter feature an extended *β*-sheet, which shares certain characteristics with the *β*-sheets found in A*β* fibrils involving a U-shaped motif [73, 74, 75, 76]. These similarities include a turn in the region E22–G29 with its center at G25 and *β*-sheets involving residues V18–E22 and G29–V40. Similar structures were also reported from NMR spectroscopic measurements for A*β*40 and A*β*42 oligomers ranging in size from tetramers to hexamers [77]. To our knowledge, a dimer structure with such a high amount of *β*-sheet and general order has never been reported before from all-atom MD simulations studying the aggregation of A*β* from disordered monomers into oligomers. We conclude that only the MD sampling of several microseconds and the usage of a force field well suited for A*β* allows the random coil to *β*-sheet transition to be simulated [78, 79]. Thus, with the current simulations we finally open the door to the nucleation process of the amyloid formation becoming observable. Using tools from chemical reaction kinetics, Linse and co-workers estimated that the average time for the appearance of the first A*β*42 oligomers that form in the lag time toward amyloid aggregation is 3 *µ*s [80]. Our simulations performed on the same time scale reveal how the structures of these oligomers evolve. Our future simulations will test whether the dimers formed in solution in the current study are indeed on-pathway toward amyloid fibril formation and whether they act as the nucleus for the subsequent amyloid aggregation process.

#### Secondary and tertiary structure

To characterize the effect of both the aggregation and membrane adsorption on the peptide secondary structure, we determined the propensity of each residue to adopt a helical conformation, to be part of a *β*-sheet or *β*-bridge, or be in a turn or bend conformation (Figure 6A). For both the dimer in solution and on the membrane considerable *β*-sheet formation is observed. With the same force field, mostly disordered conformations were sampled for A*β* during a 30 *µ*s MD simulation, with an average *β*-sheet content of about 15% [79]. This rises to ≈36% for the dimer in solution and 28% for the membrane-adsorbed dimer. As for the monomer, only little helix formation is observed. However, even if small-scale, the more apparent helix formation in the presence of the neuronal membrane is noted. This is an important observation as from NMR experiments [81, 82, 83, 84] and MD simulations using implicit membrane models [85, 86], a preference A*β* for a helix-kink/turn-helix structure, involving residues Y10–V36 with a kink or turn around residues G25–N27, in a membrane-mimicking environment such as sodium dodecyl sulphate (SDS) micelles has been reported. As in the current simulations A*β*42 mostly remained on the surface and did not noteworthy insert into it, the build-up of helix was limited. However, it is safe to conclude that the surface of the neuronal membrane inhibits *β*-sheet formation compared to the dimer in solution. This is not only evidenced by the lower amount of the *β*-sheet content for the membrane-adsorbed dimer, but also by fact that peptide 1 of this dimer, which interacts more strongly with the membrane than peptide 2, has a lower amount of *β*-sheets, but exhibits more turns, bends and random coil. On the other hand, both peptides of the dimer in solution feature a very similar secondary structure pattern along their sequence. They display a particularly high propensity for a *β*-sheet in the regions Q15–F20 of the central hydrophobic core (CHC) and A30–V40 from the C-terminal part. This excludes the residue pair G37/G38, which has a tendency for a turn formation as previously predicted by simulations and NMR spectroscopy [87].

**Figure 6:**
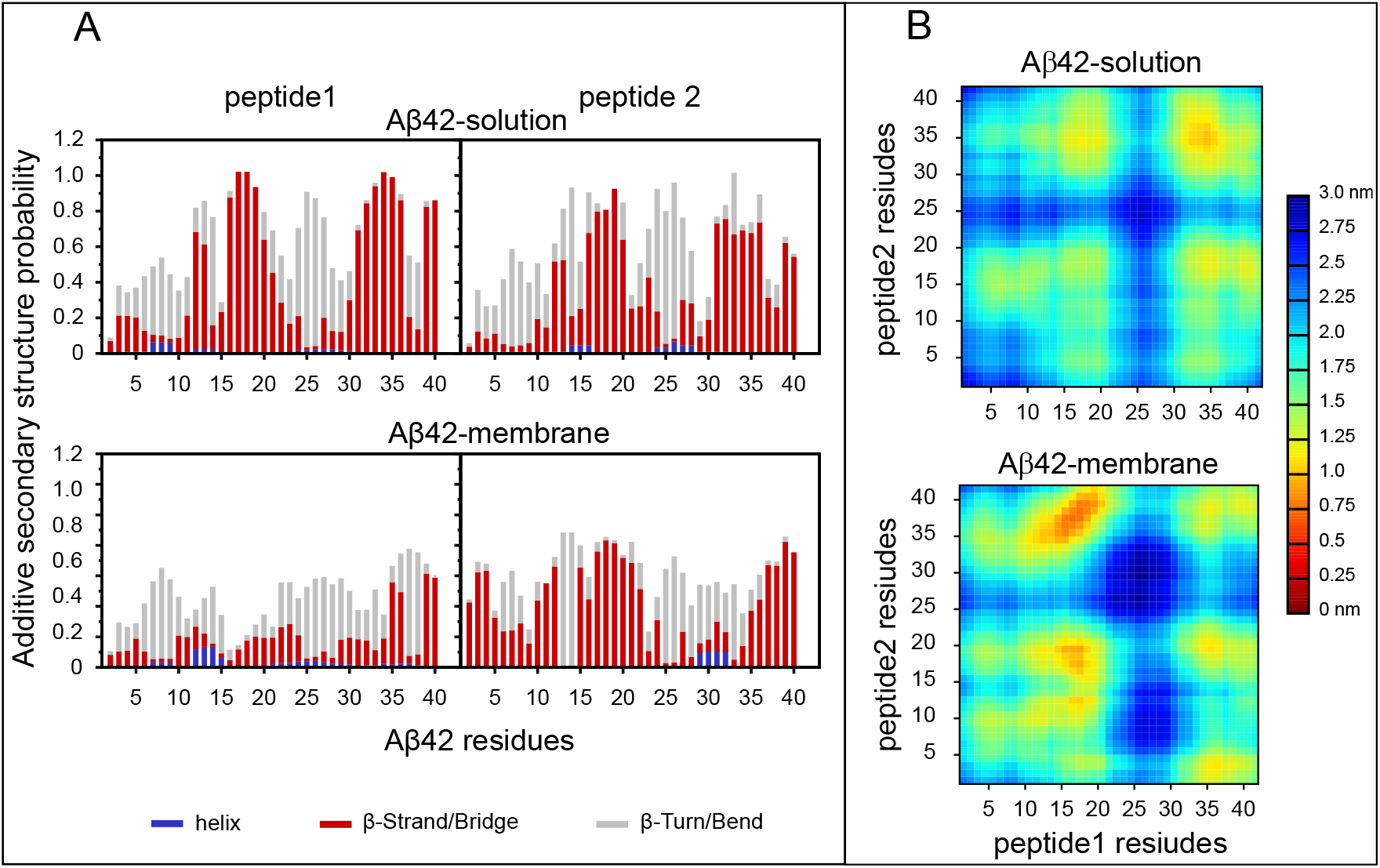
(A) Probability of secondary structure formation shown for each residue and peptide of the dimeric system in the aqueous phase (top) and in the presence of the neuronal membrane (bottom). The bars represent the additive secondary structure probabilities consisting of helix (blue), *β*-strand/bridge (red), and turn or bend (gray). The difference to 1.0 presents the random coil state. (B) The distance matrix between residues of peptide 1 and peptide 2 composing the dimer in solution (top) and in the presence the neuronal membrane (bottom). The color bar on the right indicates the average inter-residue distance (in nm)

To obtain an overview of how the different secondary structure elements are arranged with respect to each other, we calculated the distances between residues from the two peptides. The resulting distance matrices for the two dimer systems (Figure 6B) are almost symmetric with respect to their respective diagonal and are characterized by areas of high contact density (i.e., small distances) along the diagonal as well as in the upper left and lower right squares. Only the region D23–K28 does not show a noteworthy contact propensity in either dimer and for either peptide therein. This is the same region of the peptide which we identified to have a turn or bend conformation (Figure 6A). It can thus be concluded that this bend/turn region is not involved in the interpeptide interface. Instead, it gives rise to a *β*-hairpin structure for the individual peptides in the solution system (Figure 5), which in turn can assemble into interpeptide *β*-sheets. For the dimer in solution, the highest contact density is observed between the two C-terminal regions, A30–A42, which is the same region for which a high *β*-propensity was identified. These two C-terminal regions thus have a high likelihood to assemble together into a *β*-sheet in the dimer in solution. The distance matrix does not show a clear preference for either an antiparallel or parallerl *β*-sheet between these regions; therefore, both arrangements seem thus be possible. Other preferred contacts are formed between the CHC of one of the two peptides with the C-terminal region of the other peptide. Again, this must be *β*-sheet formation as these are the two regions with high *β*-propensities. The fourth area of increased contact probability is between the CHC regions of the two peptides, which for the same reason represents *β*-sheet formation. Interestingly, for the very hydrophobic regions V18-F19-F20 and I32-G33-L34 the *β*-sheet propensity reaches even values above 90%, testifying that stable *β*-sheets were formed in the solution dimer. The *β*-sheet formation coincides with regions that are involved in the cross-*β*-sheet structure found in the U-shaped A*β* fibrils [73, 74, 75, 76]. Morevoer, residues D23–K28 that we found to prefer a bend or turn conformations are the same residues which either provide the turn in the U-shaped fibril structures or the kink/turn in the helix-kink/turn-helix structures of A*β* monomers in membrane-mimicking environments [81, 82, 83, 84, 85, 86].

The distance matrix of the membrane-adsorbed dimer looks generally similar to the one of the dimer in solution. However, certain differences exist. First, the contact areas are more intense, indicating less structural diversity in the sampled dimer structures. This conclusion is supported by the reduced number of nodes in the corresponding TN (Figure 5). Second, the area without interpeptide contacts around residues D23–K28 is larger. This especially applies to peptide 1 and can be explained with the contacts that this peptide forms with the membrane instead. These residues of peptide 1 are already occupied in contacts with POPC, POPE, and GM1 (Figure S6) and are therefore not available for interpeptide contacts. Third, the order of areas with the highest contact probability or smallest inter-residue distances is different from the solution system. The smallest distances in the membrane-adsorbed dimer are observed between the CHC of peptide 1 and the C-terminal region of peptide 2. Based on the secondary structure analysis, *β*-sheet formation between these regions takes place. Furthermore, the distance matrix clearly shows a preference for an antiparallel *β*-sheet here. Another *β*-sheet is formed between the CHC regions of the two peptides. The C-terminal region of peptide 1 is, compared to the C-terminal region of peptide 2 and the dimer in solution, less involved in interpeptide contacts. Again, this is the peptide which preferentially interacts with the membrane and is therefore generally less available for interpeptide contacts. Nonetheless, some contacts of its C-terminal region with the N-terminal residues D1–5 of peptide 2 are observed. Interestingly, the latter region exhibits an increased *β*-sheet propensity, which extends up to residue Y10 (Figure 6A). Initially, it was assumed that the N-terminal region of A*β* is always disordered. However, in 2010 a simulation study had already predicted that also its first 15 residues can adopt a *β*-conformation [86], which was confirmed seven years later by cryo-electron microscopy, which was used for the determination of an A*β*42 fibril structure [88]. In the current case, *β*-sheet formation takes place between the N-terminal region of peptide 2 and the C-terminal region of peptide 1, while the first half of peptide 1 interacts with the membrane. Snapshots confirming this conclusion can be seen in Figure S5. On the other hand, also the membrane-adsorbed dimer is characterized by strong hydrophobic interactions between the two peptides, which may serve as a trap preventing more hydrophobic peptide–lipid interactions to be formed, which in turn could lead to membrane insertion that is not observed here.

#### Peptide flexibility

From the TNs but also from the contact analysis we concluded that membrane adsorption restricts the structural flexibility of the peptides. To quantify the peptide dynamics, we calculated the *S*^2^ order parameters monitoring the loss of orientation of the N–H bond vectors of the peptide backbone along with the corresponding correlation times, *τ* (Figure 7). These quantities would be directly comparable to those determined by NMR spectroscopy if the A*β*42 dimer adsorbed to a neuronal membrane were investigated with this experimental technique. Apart from a few exceptions, the dynamics of the A*β*42 residues of the dimer in solution occurs between 2 and 10 ns. The N-terminal region moves generally faster than residues from the region K16–A42. This agrees with the higher preference to form *β*-sheets in the latter region, which is also reflected by the *S*^2^ values being above 0.7 for the residues in a *β*-conformation. Such high *S*^2^ values are indicative of a folded structure, here a folded dimer. Only the turn region and some neighbored residues, ranging from E22 to A30 (and some other residues of peptide 2) have *S*^2^ values below 0.7, but above 0.5, since these are the residues which are in the turn region and therefore more mobile. The *S*^2^ values of the N-terminal residues are the smaller the closer they are to residue D1, where the order parameter has dropped below 0.3. This is in line with a disordered N-terminal region, which is confirmed by the secondary structure analysis (Figure 6) and structural snapshots (Figure5). However, the N-terminus of peptide 1 exhibited the slowest motion with *τ* ≈ 16 ns. This can only be explained by an intrapeptide contact that the N-terminus of peptide 1 must have formed for a certain time (as no marked interpeptide contact involving the N-terminus of peptide 1 is visible in Figure 6) and led to a motional restriction.

**Figure 7:**
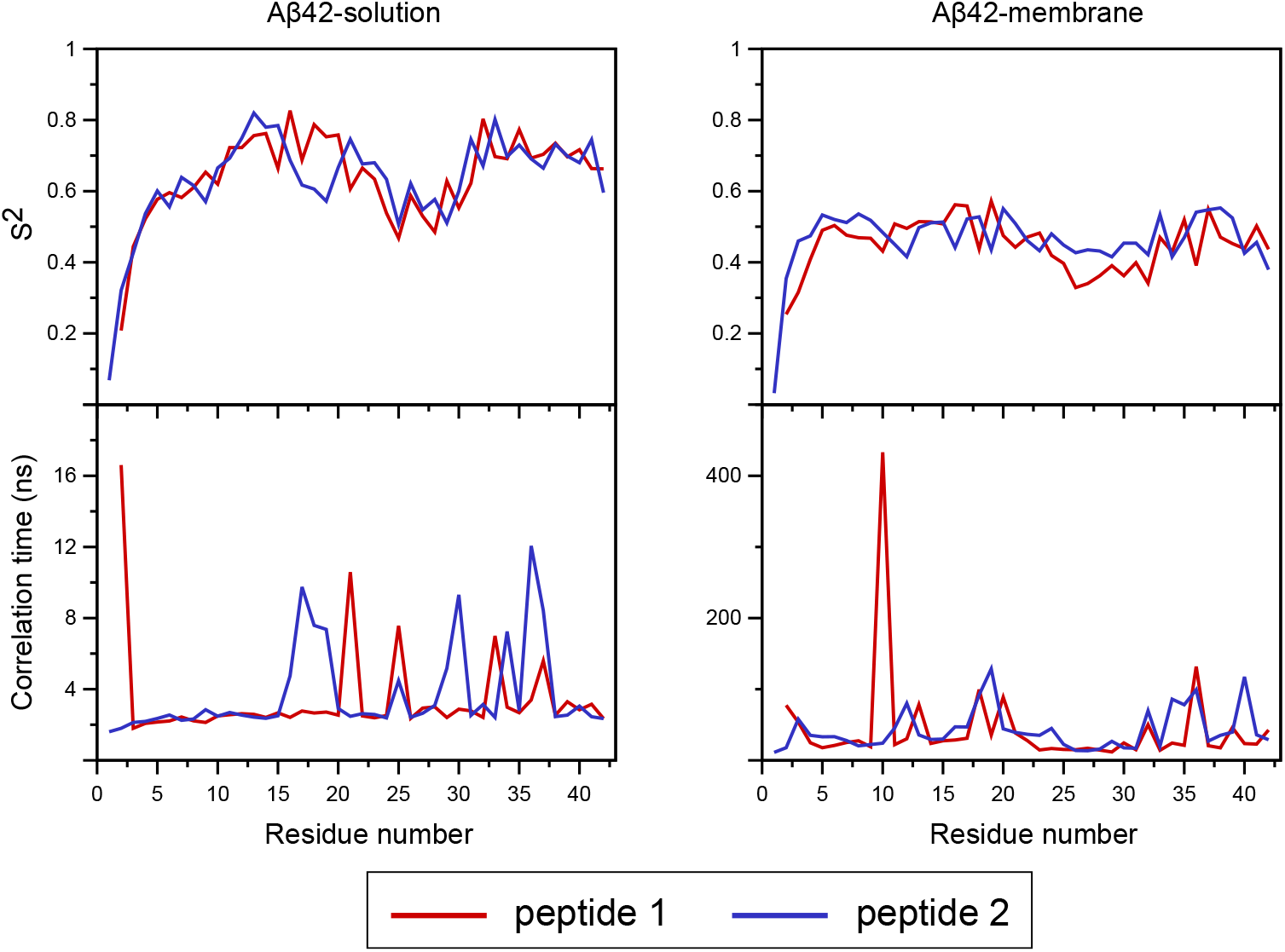
The average order parameter (top) and the correlation time (bottom) shown for each residue and peptide of the dimeric system in the aqueous phase (left) and in the presence of the neuronal membrane (right).

The dynamics of the A*β*42 dimer in the presence of the neuronal lipid bilayer is an order of magnitude slower than in solution; the slowest motion is registered for residue Y10 of peptide 1 with a correlation time of 433 ns. Interestingly, the slower motions are not accompanied by increased, but decreased order parameters in comparison to the dimer in solution; the *S*^2^ values range from 0.4 to 0.6 for most of the residues. For the N-terminal residues the order parameters are even below 0.4, but this was already the case for the dimer in solution. The overall reduction in *S*^2^ for the membrane-adsorbed dimer implies that the peptides are generally less folded than in the dimer in solution, which agrees with the reduced amount of *β*-sheet and higher propensity for random coil (Figure) 6). Thus, a picture emerges where on the one hand the overall peptide dynamics is reduced due to the adsorption on the membrane, while at the same time the interactions with the membrane reduce the local peptide order as reflected by *S*^2^.

## 3 Conclusions

In the present study, all-atom MD simulations on the microsecond time scale have been performed to elucidate the mechanism of A*β*42 dimerization in pure water and in the presence of a neuronal membrane. The consideration of a neuronal membrane consisting of six components (POPC, POPE, POPS, CHOL, SM, and GM1) is a major step forward compared to previous simulation studies on A*β*-membrane interactions, which included maximally three lipid types.

Dimerization was observed in the aqueous phase as well as on the neuronal membrane. However, the resulting dimer structures showed significant differences. Our simulations of A*β*42 dimerization in solution revealed the coil-to-*β* transition that is the first step along the amyloid aggregation pathway. The dimer conformations sampled in solution bear certain similarities with the *β*-sheets found in the U-turn shaped amyloid fibrils of A*β*. Our future simulations will unravel whether these dimers are indeed able to nucleate fibril formation. On the neuronal membrane, the dimer conformations are generally less ordered than in solution. The dimerization took place on the membrane, with one of the two peptides being preferentially adsorbed to the membrane and the other one associating with the already membrane-attached peptide without noteworthy interacting with the membrane itself. Especially the directly membrane-adsorbed peptide is characterized by a higher amount of random coil and less *β*-sheet than the peptides in the dimer in solution. The membrane adsorption is driven by electrostatic interactions between the N-terminal residues of A*β*, especially the charged ones, and the headgroups of POPC, POPE, and POPS as well as hydrogen-bonding between the sugar moieties of the GM1 lipids and A*β*42 residues across its whole sequence. GM1 is found to form clusters within the neuronal membrane, which are the preferable site for A*β* binding to the membrane surface. No insertion of the peptides into the hydrophobic region of the membrane was observed in our simulations. Instead, the interactions with the membrane rigidified both peptides, restricting their conformational diversity as the comparison to the A*β*42 dimer simulated in the aqueous phase revealed. Not only the number of conformational states is reduced as deduced from the transition networks, also the peptide motions are slowed down as revealed by the analysis of the correlation times of the N–H bond vector motions. However, while the membrane was found to have profound effects on the A*β*42 dimer, the membrane was only marginally affected by the adsorption of the A*β*42 dimer.

Our observations are in agreement with various experimental findings. Of special note is a study that analyzed the effects of glucose on A*β*42 aggregation [89]. In this study, Kedia et al. found that A*β*42 forms low-molecular weight oligomers in the presence of sugars and that these oligomers do not adopt a *β*-sheet structure. This agrees to our observation that A*β*42 dimers that preferentially interact with the sugar groups of GM1 form less *β*-sheet than the A*β*42 dimers that formed in solution. We are aware of studies by Matsuzaki and co-workers that concluded that GM1 exhibits a strong Abeta fibril seeding potential following the formation of *β*-sheet rich oligomers on GM1 clusters [90, 91]. However, these clusters are much larger than those formed in our simulations as Matsuzaki and co-workers employed ganglioside-rich (>20 mol % vs. 4 mol % in our study) membranes, where GM1 forms an interconnected network of micrometer size yielding sugar platforms in liquid-ordered membranes. As elaborated by Hof and co-workers, the scenarios for membranes with high and low GM1 contents are not necessarily contradicting each other but rather complementary [31].

Another finding by the study of Kedia et al. was that the unstructured A*β*42 oligomers that formed in the presence of glucose are able to interact with membrane bilayers [89]. Their diffusion decreased by a factor of about four upon membrane adsorption, which agrees nicely to our observation that the peptide dynamics is reduced as a result of membrane interactions. Moreover, no incorporation of the unstructured Abeta42 oligomers into the membrane was recorded [89]., which is again in agreement with our findings. However, the glucose-induced Abeta42 oligomers were able to internalize into neuronal model cells, especially with increasing glucose levels [89]. Nonetheless, it remained unanswered whether this cell penetration was a result of direct or indirect cellular uptake. Numerous studies using high levels of synthetic A*β*40 or A*β*42 indicated that such preparations are not only able to bind to membranes, yet also to perturb their structure or even create actual holes in the membrane, giving rise to ion conductance and thus leading to cytotoxicity [92, 42, 93]. However, from studies investigating A*β* in more in vivo-relevant situations, little evidence was found that major membrane disruption occurs upon exposure of neurons to natural oligomers of secreted A*β* occurring at nanomolar concentrations [94]. Instead, it is suggested that A*β*-induced neuronal dysfunction results from more subtle but sustained effects of A*β* oligomers on membrane lipids [95].

Our future simulations will show which of these two scenarios of membrane disturbance – penetration or subtle but chronic disturbance – seems more likely. To this end, even longer simulations and of larger A*β* oligomers in the presence of the neuronal membrane will be conducted. The findings from the current study support the subtle but sustained disturbance model. Moreover, if a *β*-sheet structure should be required for membrane insertion of Abeta aggregates to occur, we could even conclude that GM1 in the neuronal membrane has a neuroprotective effect as it breaks the *β*-sheet structure in the A*β* dimer. This finding would be in agreement with the neuroprotective and neurogenerative effects reported for GM1 [96, 97, 98] and the conclusion that the neurotoxic activity of amyloid oligomers increases with their *β*-sheet content [99].

## 4 Material and Methods

### Setup of the simulated systems

The systems modeled are composed of two A*β*42 peptides, which were simulated in the aqueous phase and in the presence of the neuronal lipid membrane. The initial A*β*42 structures were taken from the most populated clusters from a simulation of monomeric A*β*42 in solution [100]. The neuronal membrane model composed of 152 POPC, 96 POPE, 20 POPS, 80 CHOL, 36 SM, and 16 GM1 molecules was generated as symmetric lipid bilayer using the CHARMM-GUI interface [101]. The simulated membrane system also contained TIP3P water layers above the upper and beneath the lower membrane leaflet with sodium and chloride ions at physiological concentration of 150 mM. The two A*β*42 peptides were placed in the upper water layer at a distance of ≈ 2 nm from the equilibrated lipid bilayer surface and at a distance of *>* 1 nm between the closest atoms from the two peptides. All distances from the peptides to any of the simulation box edges were large enough to avoid interactions between the peptides with their periodic images. The total number of atoms in the modeled membrane system was ≈160,000 atoms and the box size was about 9.6 × 9.6 × 13.6 nm. The setup of the system in the aqueous phase was similar, but without a lipid bilayer, resulting in a system size of about 9.2 × 9.2 × 6.5 nm and ≈54,760 atoms.

### MD simulation conditions

The all-atom MD simulations were performed using GROMACS/2018.2 [102, 103] along with the CHARMM36m force field for A*β*42 [104] and Charmm36 for the lipids [105]. Each system was first energy minimized using the steepest decent algorithm to remove atomic clashes. This was followed by equilibration under *NV T* condition where a temperature of 310 K was regulated with the velocity-rescale thermostat [106]. Next, the system was equilibrated under *NpT* conditions to obtain a pressure of 1.0 bar. where the pressure was regulated using semi-isotropic Parrinello-Rahman pressure coupling scheme [107]. Periodic boundary conditions were set in all directions. Both the van der Waals and Coulomb force cutoffs were set to 1.2 nm in real space. The particle Mesh Ewald (PME) method was applied for calculating the electrostatic interactions. For both systems, the initial simulation was run for 2 *µ*s, from which different snapshots were randomly selected and used as starting structures for the next 5 × 2 *µ*s simulations. For the subsequent analysis, we combined the data from the six independent simulations and derived the results presented in this study.

### Analysis of the lipid bilayer properties

To evaluate the neuronal lipid bilayer properties we used different measures. The order parameter of the lipid acyl chain (*S*_CH_) is largely used to define the order of the lipid acyl chains. For this analysis, one need to define the C–H bond vectors in the required lipids and then calculates the orientation of these vectors with respect to the bilayer normal (the *z*-axis) using

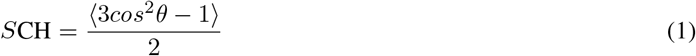

where *θ* is the angel between the C–H bond vector and the bilayer normal. The angular brackets indicate the ensemble average. The mass density profiles were calculated using the ‘gmx density’ tool along the bilayer normal. The distance between the peaks of the total density gives an estimation of the bilayer thickness. Furthermore, the bilayer thickness was calculated as the *z*-position difference between the P atoms of the lipid headgroups in the upper and lower leaflets using ‘gmx distance’ tool. The radial distribution function (RDF) provides information about the probability of finding a particle at a certain distance from another particle. We calculated the radial distribution functions of different lipid pairs in 2D (the *xy*-plane) using the ‘gmx rdf’ tool. The hydrogen bonds between different lipid pairs were determined using ‘gmx hbond’. A hydrogen bond was recorded when the angle between the donor and acceptor bonded hydrogen was between 150 and 180 degrees and the distance between the two atoms was within 0.35 nm.

### Analysis of A*β*42 properties

The order parameter *S*^2^ of a protein is an important measure that enables investigating the protein dynamics at the residue level. We used the MOPS2 (Molecular Order Parameter S2) software developed in [108] to calculate *S*^2^ for A*β*42 form the N–H bond vector autocorrelation function. The internal correlation function *C*_int_ was used after removing the overall peptide rotations. To facilitate the calculation, each trajectory was divided into subtrajectories of *t*_sub_ = 10 ns length for the solution system and 20 ns length for the membrane system. For each of the subtrajectories the order parameters were then calculated as:

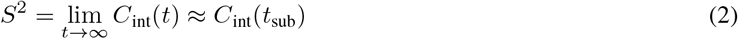

and subsequently averaged over all subtrajectories. The secondary structure of each A*β*42 residue was determined using the ‘define secondary structure program’ (DSSP) [109] invoked via the GROMACS tool ‘do dssp’. To facilitate a clear representation, the data of similar secondary structures are grouped together; *β*-strand and *β*-bridge are combined as *β*-strand/bridge, *β*-turn and bend as turn/bend, and the helix includes *α, π* and 3_10_ helices.

For the generation of the transition network (TNs) we used the ATRANET (Automated TRAnsition NETwork) software. It is a Python script available at https://github.com/strodel-group/ATRANET developed by our group to study the assembly of peptides into oligomers based on MD simulation [51]. It defines the oligomerization state by a number of descriptors depending on the properties of interest. In our case, three descriptors are used: the oligomer size, which can be 1 in the case of monomer or 2 in the case of a dimer. To define a dimer, the minimum distance between any atom of peptide 1 and any atom of peptide 2 along with the requirement of this distance to be within 0.45 nm was used. The second descriptor, the number of hydrophobic contacts between both peptides, counts the possible interpeptide atom pairs formed between the hydrophobic amino acids of A*β*42 that are within a certain cutoff (also 0.45 nm). The last descriptor is the number of residues in *β*-strand conformation, which is evaluated using DSSP and averaged over both peptides. Feeding these descriptors to ATRANET leads to a transition matrix that can be visualized using Gephi [110]. Snapshots of the representative structures from the transition network were rendered using the VMD program [111].

### Calculation of A*β*42-bilayer interactions

The peptide–lipid interactions were analyzed by calculating the interaction energy between each A*β*42 residue and the headgroup of each lipid component using ‘gmx energy’. The ‘gmx mindist’ program was employed to determine the number of contacts between each A*β*42 residue and each lipid component in the neuronal membrane. A contact was recorded when the distance between any two non-hydrogen atoms from the residue and lipid in question was within 0.5 nm. The hydrogen bond propensity was determined by the number of times a hydrogen bond was formed between hydrogen bond donating and accepting atoms in lipid pairs.

## Acknowledgments

The authors thank Dr. Michael Owen for fruitful discussions. They acknowledge funding for this project from the Palestinian-German Science Bridge financed by the German Federal Ministry of Education and Research (BMBF). B.S. received funding for this project from the Deutsche Forschungsgemeinschaft (DFG, German Research Foundation, http://www.dfg.de/) through grant number 267205415 (CRC 1208, project A07). Moreover, the computing time granted through JARA-HPC (projects JICS6C and AMYLOID-MSM) on the supercomputer JURECA at Forschungszentrum Jülich is gratefully acknowledged.

