## Supporting Information for "Amyloid-*β* peptide dimers undergo a random coil to *β*-sheet transition in the aqueous phase but not at the neuronal membrane"

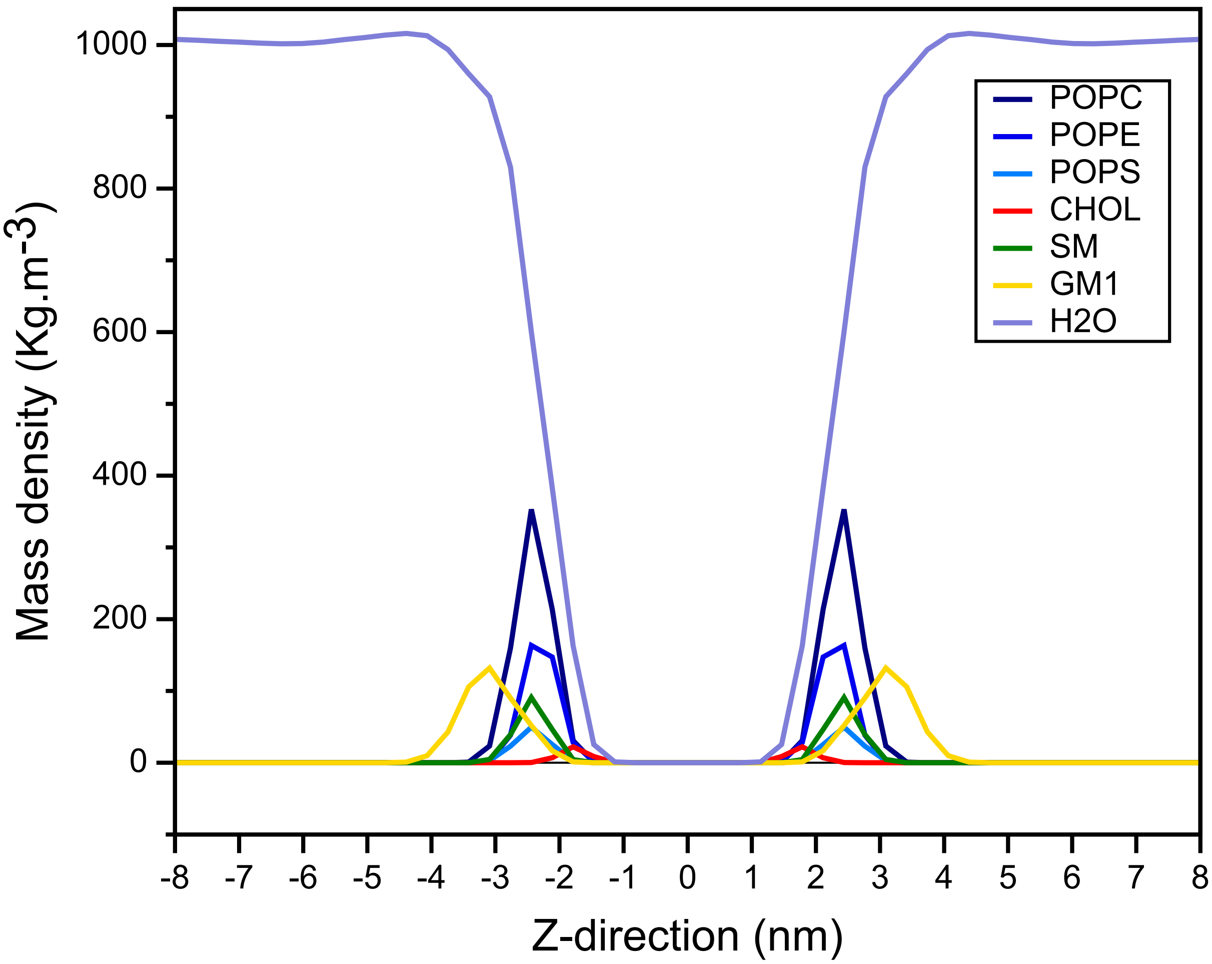


Figure S1: The average mass density profile of lipids and water along the $z$-direction corresponding to the bilayer normal. Colors are chosen according to the legend given on the right.

T

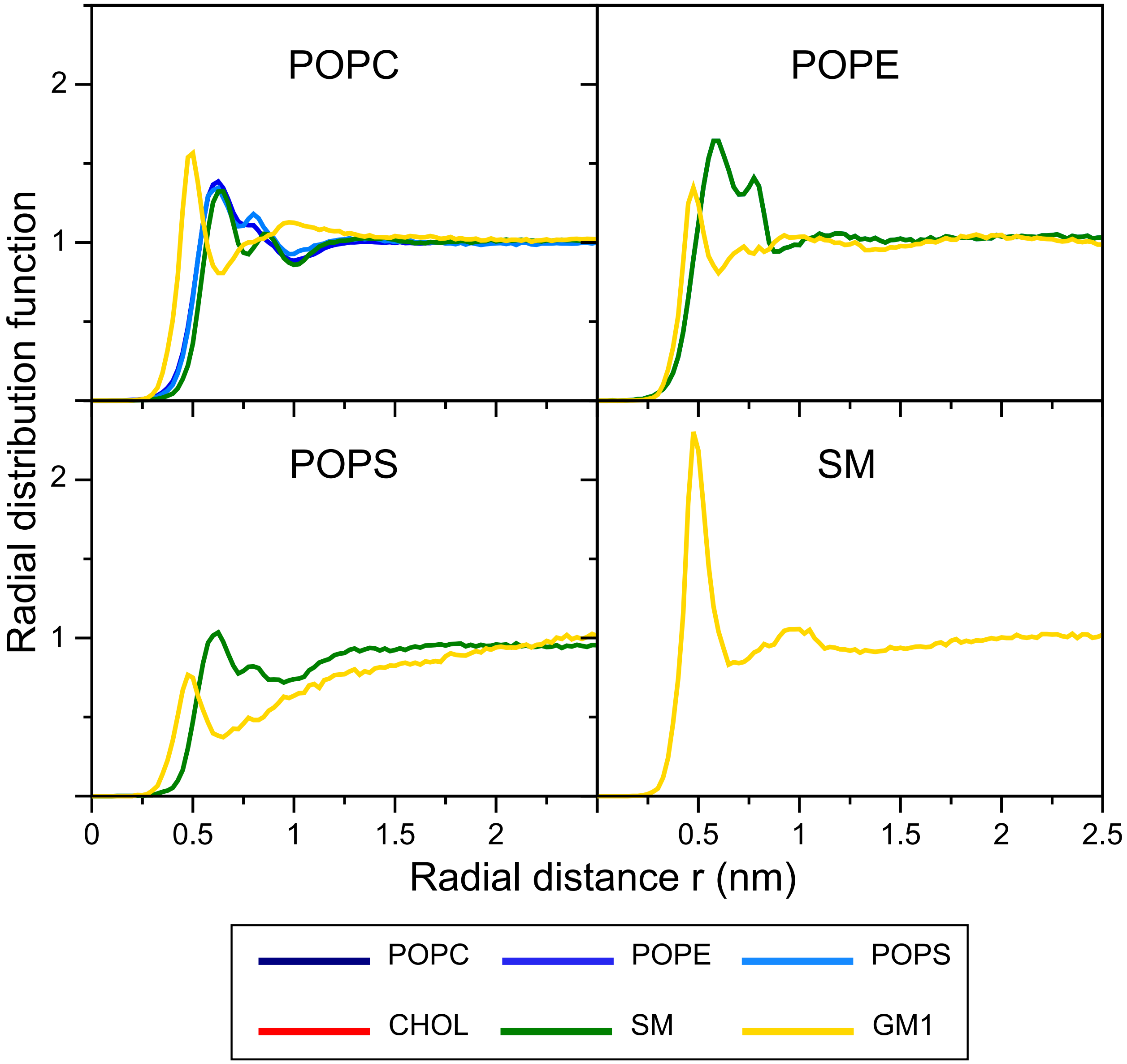


Figure S2: Radial distribution functions for mixed lipid pairings: (top left) POPC–lipid pairs; (to pright) POPE–lipid pairs; (bottom left) POPS–lipid pairs; (bottom right) SM–lipid pairings). Repeated lipid pairings, which would occur in when going from left to right and from top to bottom, are not shown. Pairings between identical lipid types are shown in the main text in Figure 2A and pairing involving CHOL are presented in Figure 2B. The P atoms of POPC, POPE, POPS, and SM and the O atoms of CHOL and GM1 were used for the RDF calculations. The $x$-axes show the distances between the respective atom pairs. The colors of the graphs refer to the lipids as indicated in the color key below the plots. Pairs with RDF > 1 are considered to cluster.

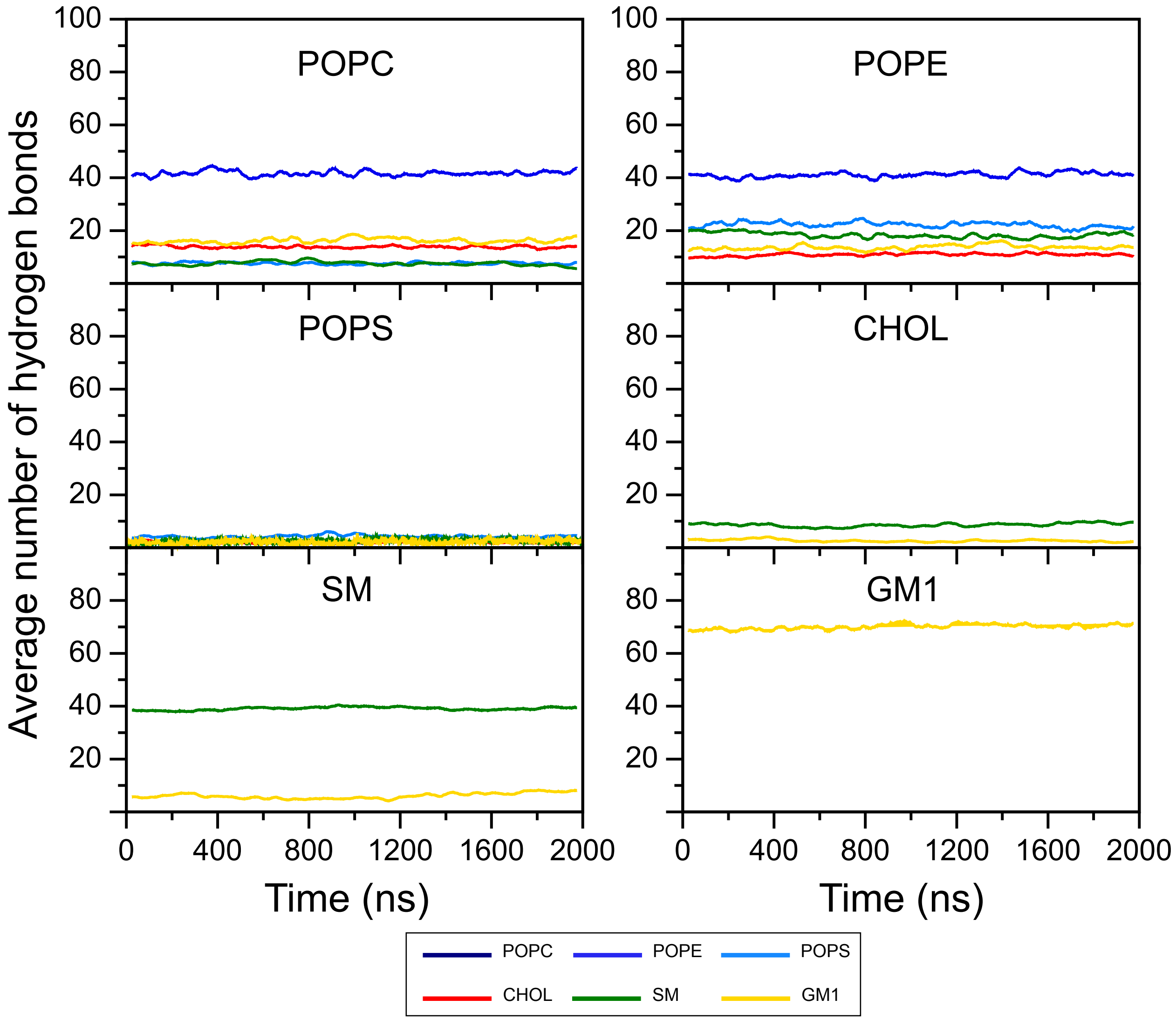


Figure S3: The average number of hydrogen bonds between different lipid pairs. Repeated pairs, which would occur in when going from left to right and from top to bottom, are not shown. The colors of the graphs refer to the lipids as indicated in the color key below the plots.

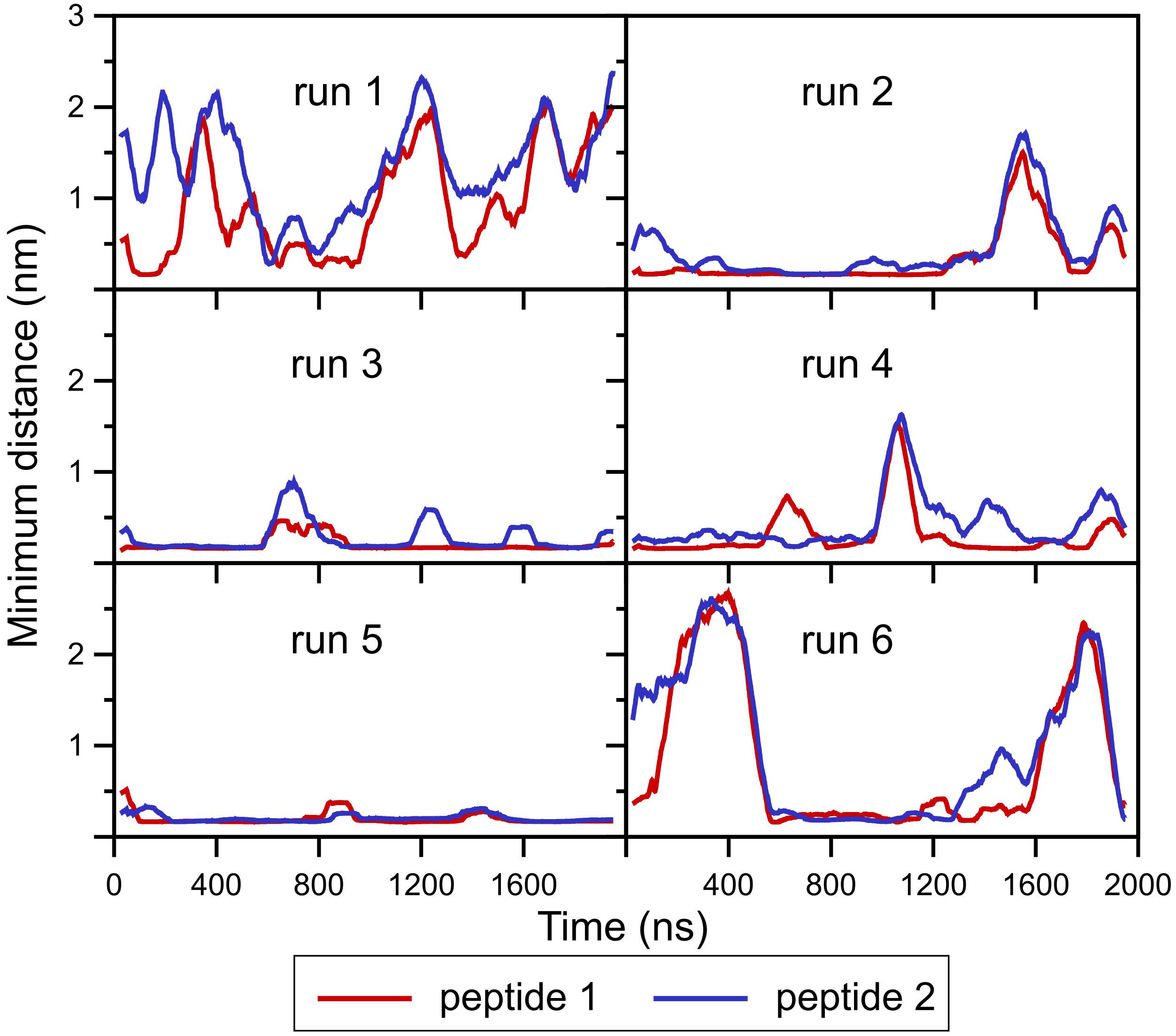


Figure S4: The minimal distance between the A$\beta$42 peptides and the neuronal membrane surface for each of the six simulations (run1–run6). Results for peptide 1 and peptide 2 are shown in red and blue, respectively.

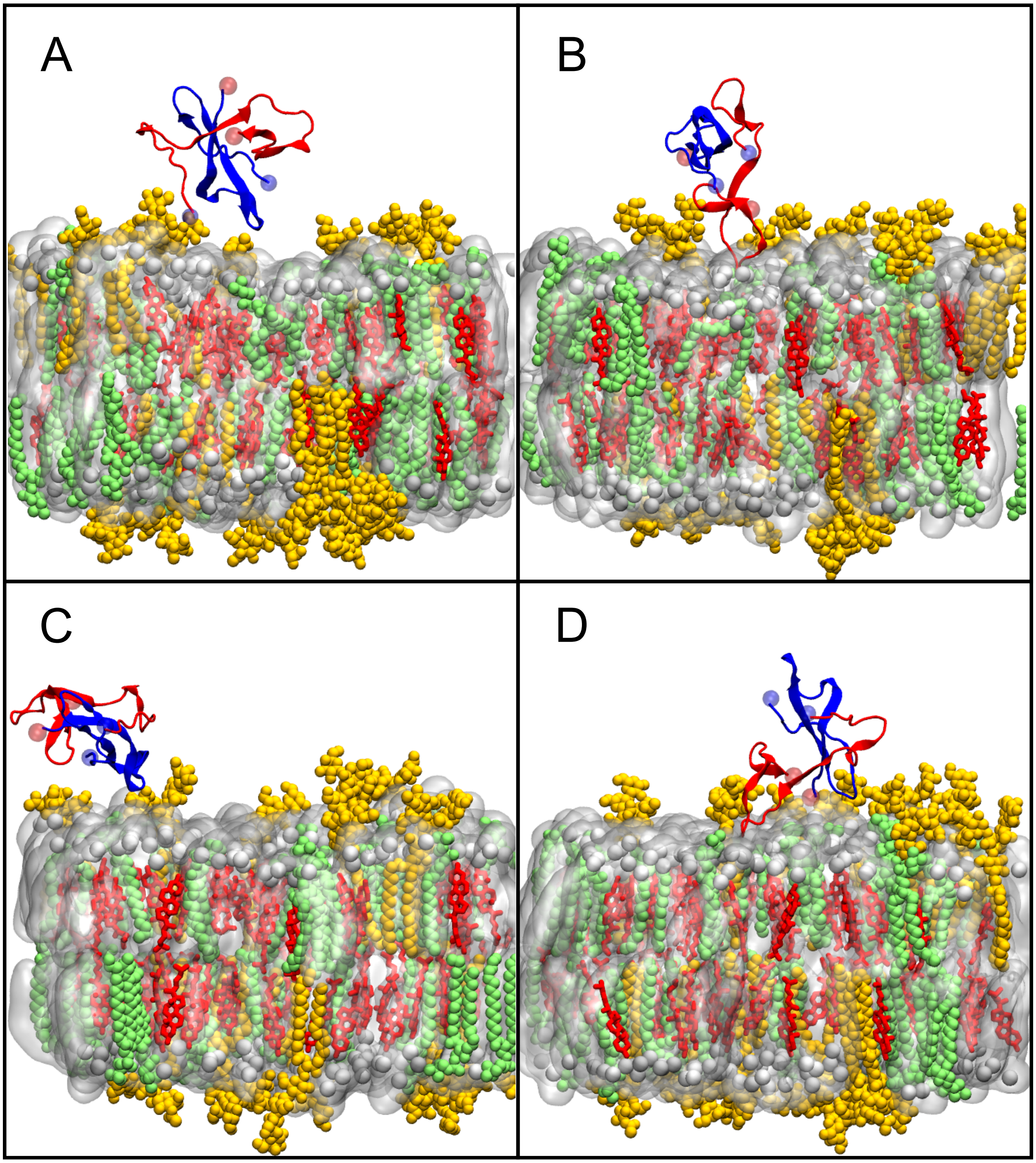


Figure S5: Snapshots of A$\beta$42 interacting with the neuronal membrane. Peptide 1 and peptide 2 are shown as cartoon in red and blue, respectively, with their N- and C-termini shown as light blue and light red spheres, respectively. POPC, POPE, and POPS of the neuronal membrane are collectively shown as silver surface with their phosphorous atoms indicated by silver spheres, CHOL is shown with red sticks, SM with green spheres, and GM1 with yellow spheres. Representative interaction patterns are provided: (A) both peptides being loosely attached to the bilayer surface, (B) peptide 1 being in close interaction with the membrane and peptide 2 being bound to peptide 1, (C) the opposite situation as in (B), (D) both peptides being in close contact with the membrane.

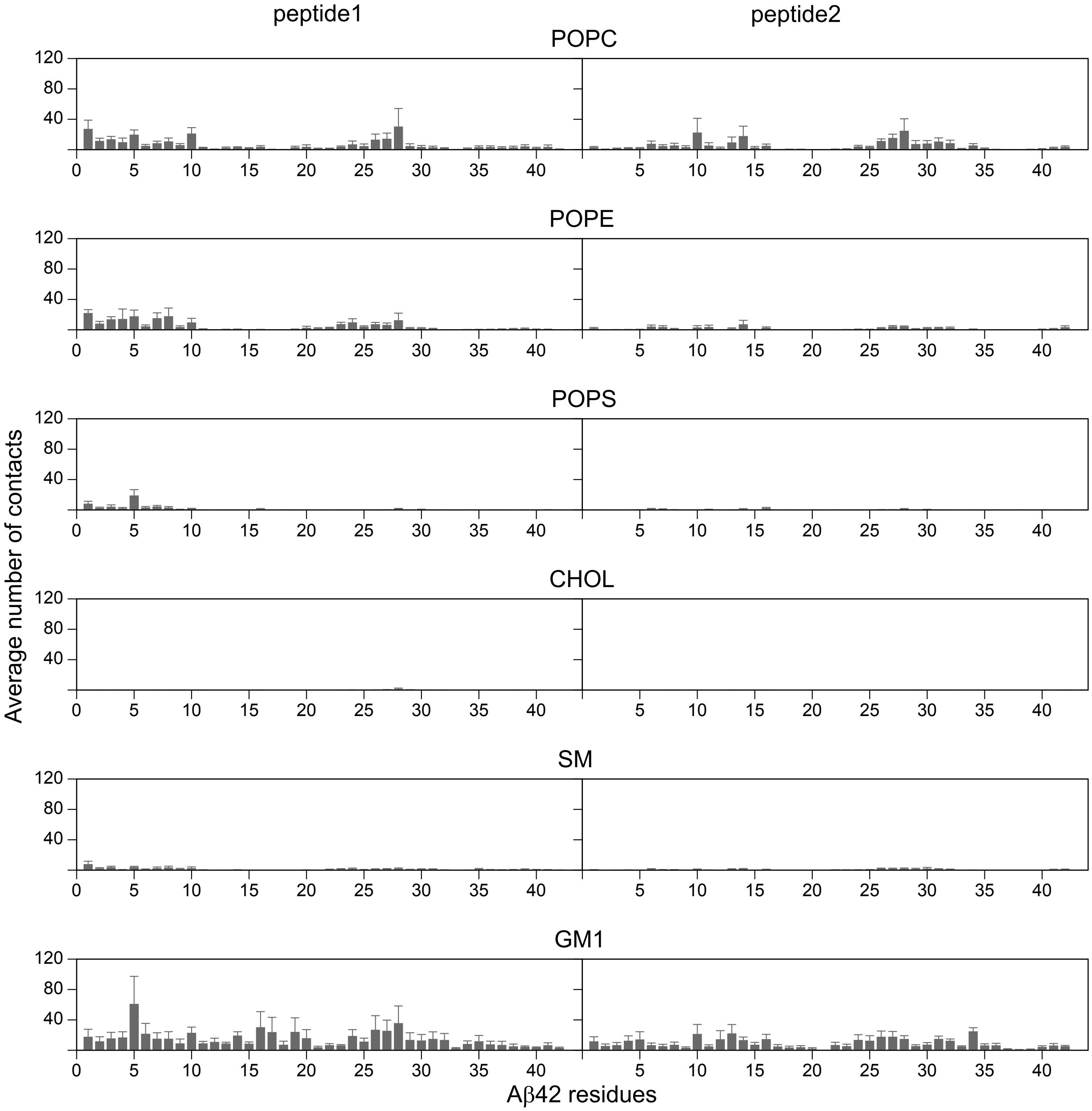


Figure S6: The average number of A$\beta$42–lipid contacts (and standard error of the mean) calculated for peptide 1 (left) and peptide 2 (right) with each of the components of the neuronal membrane (lipid names shown above the panels).

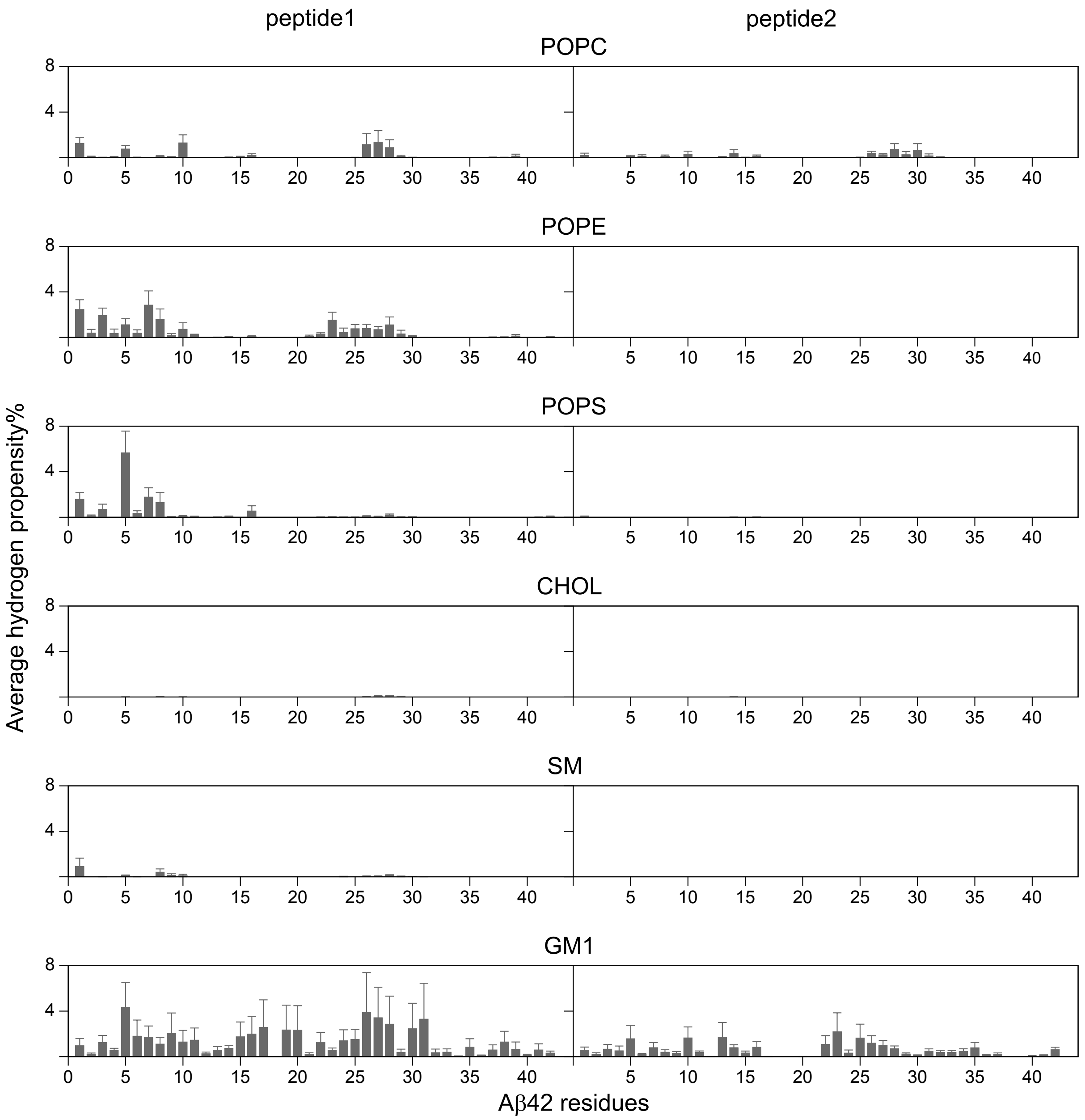


Figure S7: The average hydrogen bond propensity between A$\beta$42 and lipids (and standard error of the mean) calculated for peptide 1 (left) and peptide 2 (right) with each of the components of the neuronal membrane (lipid names shown above the panels).

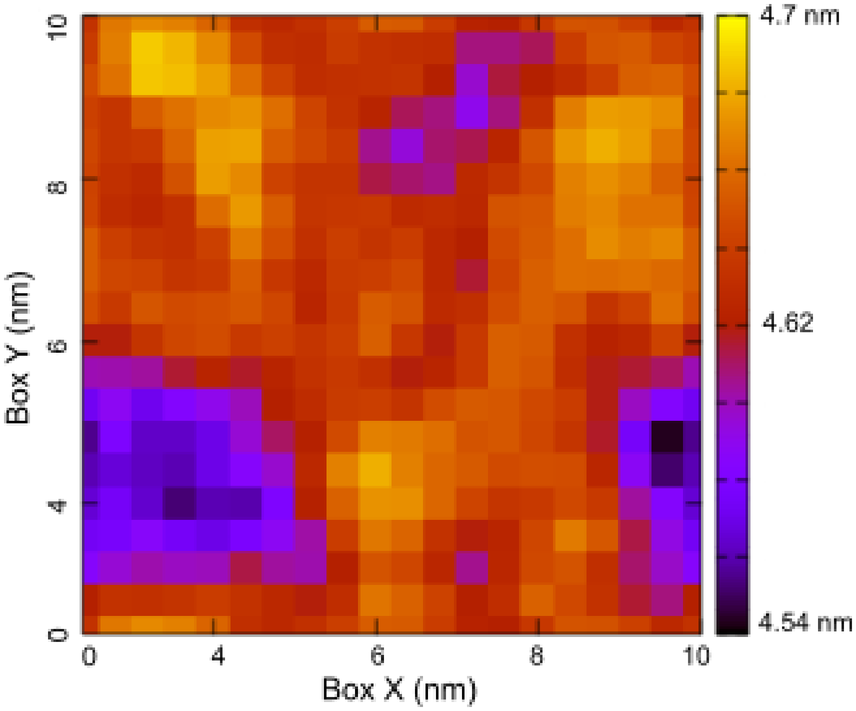


Figure S8: Average bilayer thickness calculated when the protein is within 0.5 nm of the membrane. The $x$ and the $y$-axes represent the unit cell dimension in nm. The color bar shows the thickness range in nm.
